# Calculating the maximum capacity of intracellular transport vesicles

**DOI:** 10.1101/555813

**Authors:** Erick Martins Ratamero, Stephen J. Royle

## Abstract

In eukaryotic cells, vesicles transport protein cargo to various destinations in the cell. As part of an effort to count the number of cargo molecules in vesicles in cells, we asked a simple question: what is the maximum number of cargo molecules that can be packed into a vesicle? The answer to this question lies in the Tammes Problem, which seeks to determine the packing of circles on a spherical surface. The solution to this problem depends on the sizes of the vesicle and the cargo. We present here the computational results that determine the maximum theoretical capacity of a range of biologically relevant cargo in a variety of meaningful vesicle sizes. Our results could be generalized in order to describe an equation that allows the calculation of the approximate maximum capacity of any vesicle, whatever the size; provided that the size and geometry of the cargo is known.

## Introduction

Inside all eukaryotic cells, proteins and lipids are moved around by vesicle transport. To move from one membrane compartment to another, a protein must be packaged into a vesicle where it travels as ‘cargo’ until the vesicle fuses at the destination membrane. Vesicle transport controls 1) the secretion of newly synthesised proteins from the endoplasmic reticulum (ER) to the Golgi apparatus, through the Golgi and on to the cell surface; 2) the endocytosis of receptors from the plasma membrane and their subsequent degradation or recycling back to the plasma membrane. Since the identity and function of each membrane domain is determined by its protein and lipid composition, vesicle transport affects virtually all cellular processes including signalling, migration and nutrition (Bonifacino and Glick, 2004).

Intracellular transport vesicles are approximately 30 nm to 100 nm in diameter. The vesicles within a clathrin cage are typically 40 nm to 100 nm (Vigers et al., 1986). Transport vesicles associated with the Golgi are approximately 60 nm (Cosson et al., 2002). Synaptic vesicles in neurons are 40 nm (Takamori et al., 2006). Work from our own lab has identified 30 nm diameter vesicles (Larocque et al., 2018). Outside of these ranges, there are larger, non-spherical carriers such as tubulovesicular recycling vesicles, while the lower limit for a vesicle is estimated to be 20 nm diameter (Huang, 1969).

There has been recent interest in quantifying the absolute numbers of proteins associated with intracellular vesicles. Proteomic estimates of protein composition have been obtained for COPII and COPI vesicles (Adolf et al., 2019), AP-1/clathrin-coated vesicles (Hirst et al., 2015) and for synaptic vesicles (Takamori et al., 2006; Wilhelm et al., 2014). In the latter two cases these data were used to build 3D models of an average vesicle which contains cargo. An alternative route is to directly count protein numbers at individual vesicles using light or electron mciroscopy (Cocucci et al., 2012; Grassart et al., 2014; Mund et al., 2018; Sochacki et al., 2017; Clarke and Royle, 2018). As part of our own efforts to count cargo molecules in intracellular vesicles, we wanted to calculate the maximum capacity of a given vesicle, to model the feasibility of our approach. Finding the theoretical maximum capacity of a vesicle requires solving a spherical packing problem known as the Tammes Problem (Tammes, 1930).

Our calculations describe the maximal capacities of a range of intracellular vesicle sizes and for a variety of cargo shapes. While these results are theoretical, and such density of packing is unlikely to occur under normal conditions, they highlight the enormous flux capacity of vesicular transport and provide a useful dataset to model labeling efforts in the future.

## Methods

### Packing problem

An important part of our approach is the reduction of all packing problems to the Tammes Problem, which involves packing a certain number of circles on the surface of a sphere while maximising the minimum distance between circles. The Tammes Problem was originally formulated to describe the pores on the surface of a pollen grain (Tammes, 1930). If the pores are considered as circles and the pollen grain as a perfect sphere, the problem is a specialized form of the generalized Thomson Problem. Here, *N* electrons on the surface of a sphere repel each other in order to minimize their energy configuration. In the Tammes Problem, *N* circles on the surface of a sphere are packed in order to maximize the minimum distance between them. To approach our problem from a theoretical point of view, we made a number of approximations and assumptions in order to reduce all packing problems to the Tammes Problem. First, all vesicle surfaces were approximated as perfect spheres. Second, cargo proteins were considered as either cylinders or spheres, with circular intersections with the vesicle sphere. Cylinders have their main axis aligned with radial vectors from the vesicle sphere, meaning that all protein intersections with that sphere will be circles. These approximations allow us to perform all calculations on perfect spheres, even when solving a packing problem at places other than the vesicle sphere. In this way, the sphere radius can be adjusted to account for protein extension into or away from the vesicle (Figure 1).

**Figure 1.**
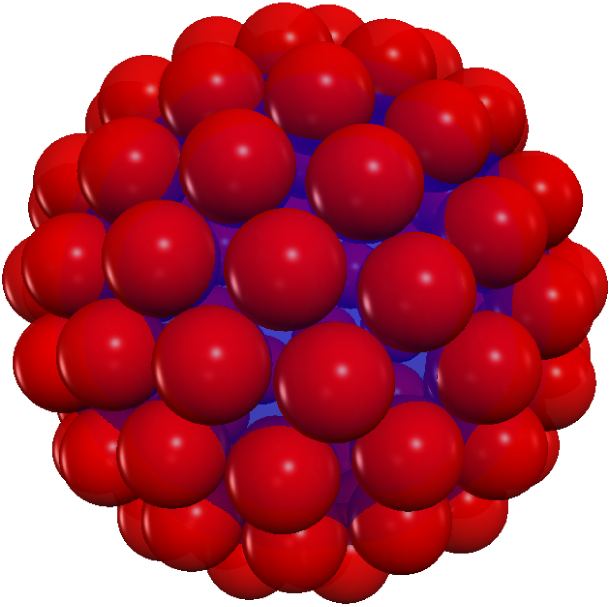
Illustration of the packing problem. Packing of 100 red spherical ‘cargo’ proteins on a blue sphere. Coordinates for the sphere centres and their radius were obtained by the approximated Tammes solver. The image shown is generated after maximizing the minimum distance between them. The blue sphere represents the sphere where packing occurs, i.e. the vesicle sphere adjusted for protein extension.

Given these approximations, we have moved the problem from packing complex protein structures on the surface of a vesicle sphere to packing circles on the surface of an equivalent sphere. Now, this problem is equivalent to maximising the minimum distance between points on a sphere, where the points are the center of each circle. Packing can be achieved as long as the smallest distance between two points is larger than the diameter of the circles to be packed. This is because the closest distance between two circles is equal to the sum of their radii, which is equal to the diameter for two circles of equal size.

This formulation of the problem relies on the local curvature of the sphere being relatively small compared to the radius of the circles, so that the euclidean distance between two centres of circles is approximately the same as the distance between them on the sphere. This is not an issue for large numbers of circles, but that approximation is weaker for very large proteins or very small vesicles.

### Computational solution

In the previous section, we transformed our problem from packing complex proteins on the surface of a vesicle into maximising the minimum distance between points on a sphere. This new problem is now equivalent to the Tammes Problem, for which a number of approximate solvers are readily available. Note that exact analytical solutions for this problem only exist for a very limited range of values (Musin and Tarasov, 2015).

We reused an approximate solver, for which the source code was readily available and easy to edit (Pruss, 2017). This solver is based on the idea of considering the points to be spread on the surface of a sphere as particles with repulsive forces between them. With that, it is relatively easy to run a computer simulation of their dynamics from a random initial position, by considering the repulsion between particles and some frictional damping.

The solver outputs the minimum distance between two points after optimising *N* point positions on the surface of a unit sphere. Therefore, given a number of points, *N*, we obtain an equivalent circle radius *r*_equiv_ that needs to be scaled by a factor equal to the actual vesicle size we are interested in. Of course, in our formulation of the problem we have the opposite: our input is a circle radius *r*, given by the protein structure, and a vesicle radius *R*; the output we want is the number of circles *N* that can be packed on the vesicle surface.

A few adaptations were therefore necessary. We start by trying to pack an *N*_0_ number of circles of radius *r*_equiv_ onto a unit sphere. The radius *r*_equiv_ is calculated by dividing the radius of the intersection between protein and sphere (vesicle or packing sphere) by the radius of the vesicle we are interested in, so that we can operate over a unit sphere instead.

Running the approximate Tammes solver with parameter *N*_0_, we get an output of *r*_0_. As long as that *r*_0_ (which is the minimum distance between the *N*_0_ points spread over a unit sphere surface) is larger than the *r*_equiv_ radius of the protein-vesicle intersection circle, we consider that we can fit *N*_0_ proteins on the surface of the vesicle.

If that packing problem is successful, we can proceed with trying the same problem with *N*_0_ + 1 proteins, then *N*_0_ + 2 proteins and so on. Eventually, we will run into the situation where the output *r*_out_ < *r*_equiv_. At that point, we cannot pack any more proteins on the surface of the sphere, and we take the last successful packing input *N*_fit_ as the maximum number of proteins.

Due to the stochasticity embedded into the approximate Tammes solver, we repeated this process ten times for each protein. Furthermore, we also varied the vesicle radius by changing the value of *r*_equiv_ accordingly. We have chosen a range of typical vesicle sizes with radii of 15 nm to 50 nm. The results across ten simulations were very consistent, with the highest variability for small cargo packing onto the largest vesicle sizes. Even here, the solutions were on the order of 1000 to 2000 circles, with a standard deviation of only 1 to 3. We therefore present the mean value from the ten simulations in the plots of specific cargo types. The minimal variability in the end allowed us to approximate a general solution to this problem.

### Protein modeling

We selected four model cargo molecules to explore the maximum capacity of intracellular vesicles. All four are commonly found on intracellular vesicles. First, vacuolar ATPase (vATPase) – a large multipass transmembrane protein complex that acidifies organelles via proton-pumping. Second, G-protein coupled receptor (GPCR) – compact seven transmembrane proteins that couple to small G-proteins. Third, Rab GT-Pase (Rab) – small proteins which bind the cytosolic face of the vesicle via lipid modification. Four, transferrin receptor (TfR) – a single-pass transmembrane protein with a large extracellular domain that binds transferrin. To model these proteins, we used the following structural models and measured their dimensions using PyMol (DeLano Scientific). The bilayer was considered to be 4 nm thick. First, yeast vATPase (PDB = 6C6L), the widest portion was 12.2 nm, the radius of the packing sphere was equal to the vesicle radius minus 2, i.e. the luminal face of the bilayer was used (Roh et al., 2018). Second, beta2-adrenergic receptor (PDB = 3SN6), the width of 4 nm, again using the luminal face of the bi-layer as the packing sphere (Rasmussen et al., 2011). Note that the only the receptor, minus the lysozyme moiety was used, ignoring the G-protein. The positioning of these two proteins was aided by their entry in OPM (Lomize et al., 2012). Third, Rab5a (PDB = 1N6H) with a width of 4.1 nm oriented according to molecular dynamics work (Zhu et al., 2003; Edler and Stein, 2017). This placed the widest portion 2 nm from the cytoplasmic surface of the vesicle, 4 nm from the middle of the bilayer. Fourth, unliganded transferrin receptor (PDB = 1CX8), the widest part of the dimer is 9.8 nm and the placement of this part of the molecule at 7.6 nm from the luminal surface of the vesicle was estimated using the liganded TfR structure (PDB = 1SUV) (Cheng et al., 2004; Lawrence et al., 1999). The respective packing sphere radii were therefore *R -* 2, *R -* 2, *R* + 4, and *R-* 9.6.

### Data and software availability

Source code and data is available at GitHub and Zenodo (Ratamero, 2019).

## Results and Discussion

### Exploring maximum vesicle capacities using four different cargo types

We used four typical cargoes to explore the maximum capacities of intracellular vesicles. These cargoes are all well-studied protein that were chosen due to their ubiquity and varied size. The dimensions of vATPase, GPCR, Rab and TfR are described in Methods and shown schematically in Figure 2A. Computer simulations for packing of these four cargo types into vesicles with radii of 15 nm to 50 nm was done in order to find the maximum capacity in each case (Figure 2B).

**Figure 2.**
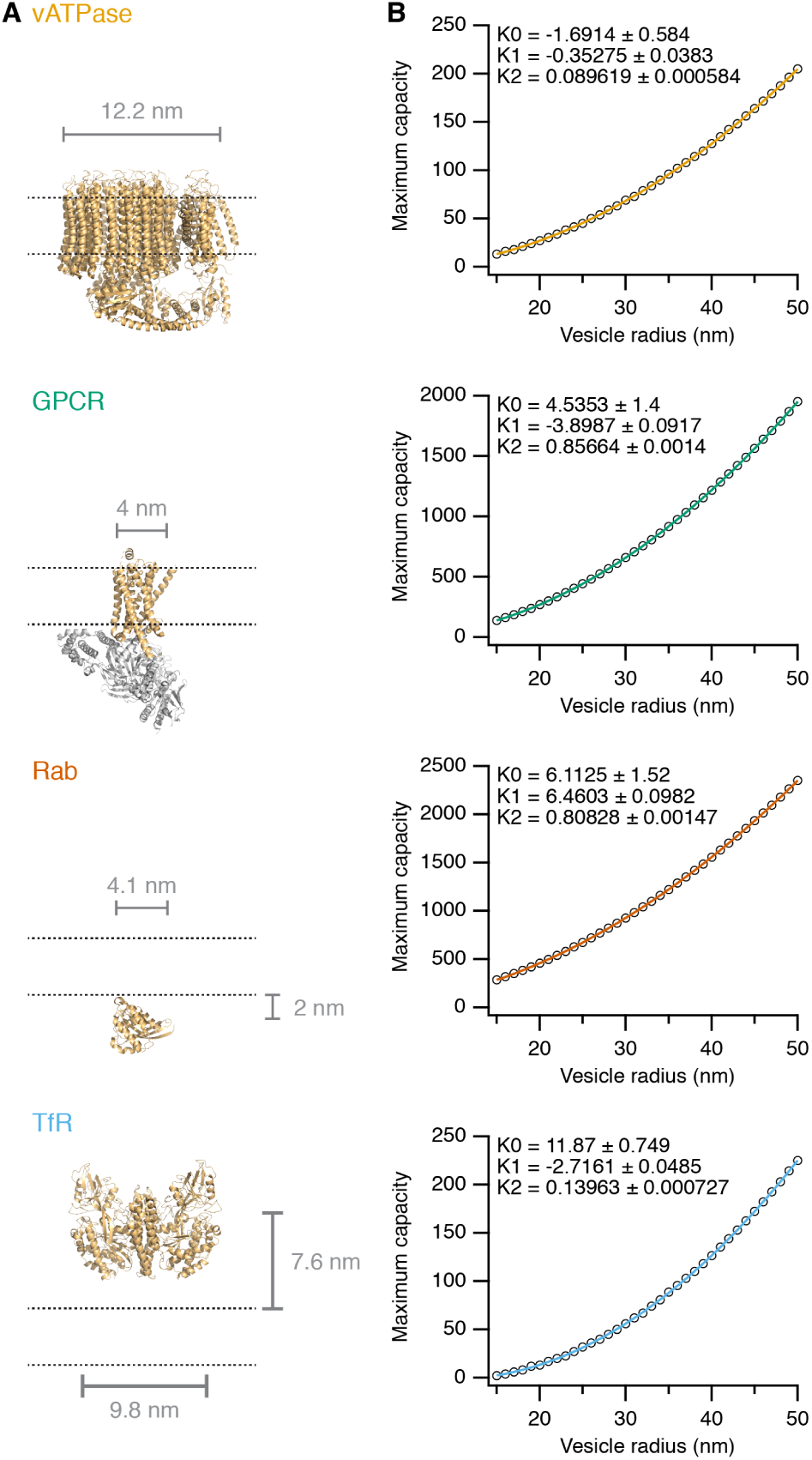
Maximum vesicle capacities calculated for four cargo types. The packing of vATPase (6C6L), GPCR (3SN6), Rab (1N6H) or TfR (1CX8) in vesicles with radii from 15 nm to 50 nm. **A**, Schematic illustrations of structural models for each of the four cargoes is shown relative to a vesicular membrane (the lumen of the vesicle is above). **B**, Plots to show the maximum capacity for each cargo at a variety of vesicle sizes. A polynomial fit is shown of the form *y* = *k*_0_ + *k*_1_ *x* + *k*_2_ *x*^2^. Coefficients and error are shown inset.

The maximum capacity of vesicles was surprisingly large in some cases. For example, a vesicle of 50 nm radius can pack almost 2500 copies of Rab5a. A vesicle of the size typically found with a clathrin coat (radius of 30 nm) could contain up to 660 copies of a GPCR. We emphasize that these values are theoretical maxima, and that they do not account for 1) steric hindrance between cargo molecules, 2) biological inefficiencies of packing, 3) mixed cargo types in vesicles. To highlight this difference, the current best estimate for Rab proteins on a synaptic vesicle is 25 copies for a vesicle with a radius of 20 nm (Takamori et al., 2006), whereas the maximum capacity is approximately 460 copies.

In each case, a polynomial function could be fitted to the simulated data (Figure 2B). A second-order polynomial was chosen since we expected the maximum capacity to roughly scale with the total vesicle surface, which in turn scales with the square of the vesicle radius. Overlay of these fits allows a direct comparison which high-lights that the maximum capacities for small cargo are much higher than wider cargo or cargo that protrudes into the lumen (Figure 3A).

**Figure 3.**
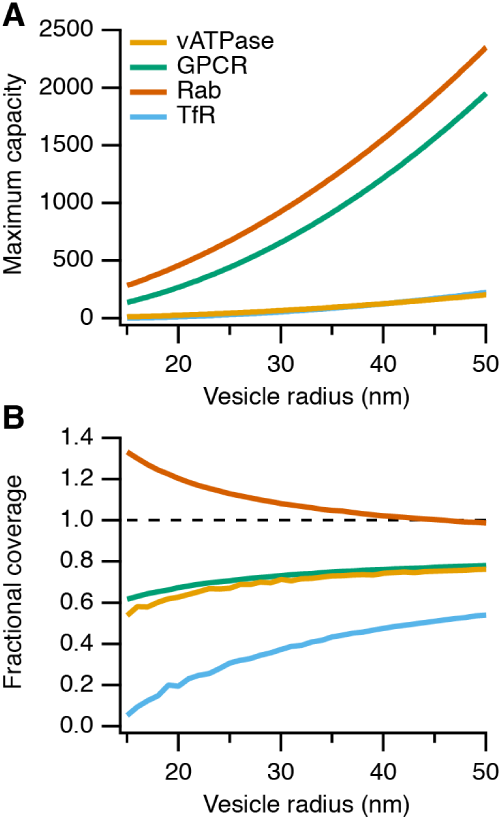
Maximum vesicle capacities and coverage for four cargo types. **A**, The fits from Figure 2 are plotted together to allow comparison between the four cargo types. **B**, Fractional coverage of the cargo types as a function of vesicle size. Fractional coverage is the total cargo area (circle area) divided by the vesicle surface area.

We next wondered what is the vesicular coverage of different cargoes at maximum capacity? To look at this, we calculated for each vesicle size the vesicle surface area and the corresponding total area of packed circles. Fractional coverage was therefore the total cargo area divided by the vesicle surface area (Figure 3B). Two interesting things about this plot are: Firstly TfR, which is a wide cargo protruding far into the vesicle lumen, is packed at low rates and results in very low fractional coverage. The cellular solution to such cargo types is to package them into larger vesicles. One example of this phenomenon is that feeding cells with EGF results in larger intracellular vesicles (Edgar et al., 2014). Secondly, supramaximal packing of Rab can be seen, i.e. the capacity of vesicles for Rab exceeds 1 at vesicle radii smaller than 45 nm. This is because the Rab is on the cytosolic face of the vesicle and as such, its packing sphere is larger than the vesicle sphere.

In summary, the cargo proteins with larger cross-sectional areas can be packed in smaller numbers than the smaller proteins. Furthermore, proteins that coat the inside surface of vesicles have decreased numbers compared to those on the outside of vesicles.

### Approximating a general solution to cargo packing

As noted in the Methods section, we were surprised – given the stochasticity involved in our computational solver – that the variability of maximum capacity was so low. This meant that, while an exact analytical solution to the Tammes Problem for all integers is not practical, we should be able to use the data from the simulations to state an approximate generalized solution for cargo packing. As described in Methods, the key parameter is the ratio between the packing sphere radius and the cargo radius. We call this parameter, *q*. Recall that the radius of the packing sphere is the vesicle radius allowing for protein extension into or away from the vesicle. Taking our simulation results for all four cargoes and plotting them as a function of *q*, reveals that all the points follow a second-order polynomial of the form *y* = *k*_0_ + *k*_1_*x* + *k*_2_*x*^2^ (Figure 4A). Therefore, the maximum capacity *C*_max_ can be calculated using *q* according to Equation 1.

**Figure 4.**
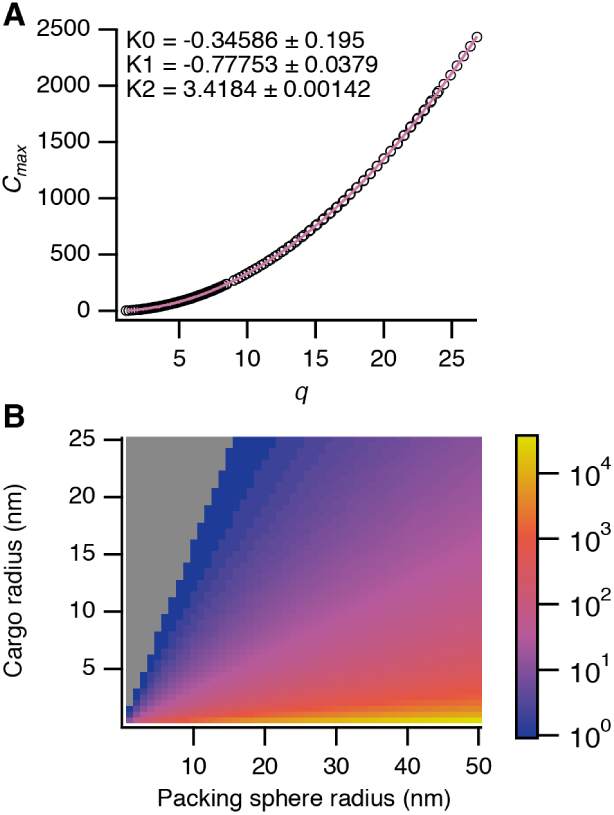
General solution to calculating the maximum capacity of intra-cellular vesicles. **A**, Maximum capacities (*C*_max_) of all four cargo types plotted as a function of *q*, the ratio of cargo radius to packing sphere radius. The grouped data were well fit by a second-order polynomial of the form *y* = *k*_0_ + *k*_1_ *x* + *k*_2_ *x*^2^. Coefficients and errors for the fit are shown inset. **B**, The fit shown in A could then be used to calculate a matrix of values for a range of cargo sizes at various packing sphere radii. In the plot, *C*_max_ is shown using ametrine look-up table. Note the log scaling, min = 1, max = 34105, gray = 0.

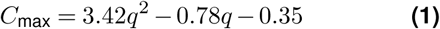

Using Equation 1, we could then produce a matrix of values for different cargo sizes and various packing sphere sizes for illustrative purposes (Figure 4B).

### Conclusion

In this work, we showed how the the problem of calculating the maximum capacity of intracellular vesicles can be reduced to the Tammes Problem, wherein a number of circles is packed on the surface of a sphere. Using a computational solver we calculated the maximum theoretical capacities for proteins on the surface of vesicles typical of the sizes encountered in cells. This allowed us to present a generalized solution to calculate the maximum capacity of a vesicle without running the solver. We hope the information in this paper will be useful when thinking quantitatively about vesicle transport in cells.

## ACKNOWLEDGEMENTS

The authors gratefully acknowledge CAMDU (Computing and Advanced Microscopy Unit) for their support and assistance in this work.

## AUTHOR CONTRIBUTIONS

Conceptualization, E.M.R. and S.J.R.; Methodology, E.M.R. and S.J.R.; Software, E.M.R.; Investigation, E.M.R.; Writing and Original Draft, E.M.R. and S.J.R.; Review and Editing, E.M.R. and S.J.R.; Funding Acquisition, S.J.R.; Resources, S.J.R.; Supervision, S.J.R.

